# Exosome-mediated LNA anti-miR delivery achieves potent miR-21 inhibition and PTEN restoration in podocytes

**DOI:** 10.64898/2026.07.17.738885

**Authors:** Tim Lange, Joanne Ern Chi Soh, Lea-Katharina Friedrich, Paul Luca Götsch, Claudia Weber, Doreen Biedenweg, Rabea Schlüter, Una Janke, Mihaela Delcea, Nicole Endlich

## Abstract

**Background:** Podocyte injury and loss are central drivers of chronic kidney disease (CKD) progression and are involved in the majority of glomerular diseases. Among the molecular pathways implicated in podocyte dysfunction, microRNA-21 (miR-21) is consistently upregulated during glomerular injury and suppresses protective target genes including PTEN. Although anti-miR-21 approaches have shown high efficacy in preclinical models for Alport Syndrome and diabetic nephropathy, clinical translation of free anti-miR-21 oligonucleotides remains challenging, potentially reflecting limited glomerular and podocyte target engagement. Here, we show that an exosome-mediated delivery of an LNA-miR-21 inhibitor results in a higher intracellular uptake, leading to functional miR-21 inhibition and PTEN restoration in podocytes.

**Methods:** Exosomes were isolated from immortalized murine podocytes SVI and were directly loaded with fluorescently-labeled LNA anti-miR-21 (LNA-21) and anti-miR-Ctrl. Exosome integrity was assessed by transmission electron microscopy, dynamic light scattering and Western blot analysis of CD9 and TSG101. Uptake efficiency and intracellular delivery were analyzed by confocal laser-scanning microscopy, time-series imaging (2-48 h) and (imaging) flow cytometry. Functional efficacy was evaluated by TaqMan RT-qPCR and PTEN protein expression in differentiated SVI podocytes, primary murine podocytes and puromycin aminonucleoside (PAN)-injured primary glomeruli.

**Results:** Exosome number, morphology and marker expression remained unchanged after direct LNA-miR-21 loading. Time-series imaging demonstrated progressive intracellular accumulation of exosome-delivered LNA-21 cargo from 2 to 48 h, whereas free LNA-21 showed no detectable uptake. Flow cytometry and imaging flow cytometry confirmed an efficient (76.6%) intracellular delivery exclusively via exosomes. Exosome-mediated LNA-21 delivery induced a robust miR-21 suppression, resulting in an 99% reduction in immortalized podocytes, a 97% reduction in primary podocytes and a 97% reduction in PAN-injured glomeruli (all p < 0.05). In contrast, free LNA-21 did not achieve significant suppression in all investigated models. To evaluate the functional consequences of miR-21 inhibition, we examined PTEN expression, as PTEN is a known direct target of miR-21. Notably, restoration of PTEN protein expression was observed only following exosome-mediated LNA delivery which was confirmed by Western blot and immunofluorescence, suggesting efficient functional inhibition of miR-21 in podocytes. Transfection of scrambled controls confirmed sequence specificity throughout.

**Conclusion:** These findings demonstrate that exosome-mediated LNA delivery overcomes the limited uptake of free inhibitor oligonucleotides in podocytes and enables functional inhibition of miR-21 across multiple experimental systems. These data provide mechanistic proof-of-principle for exosome-based miR inhibitor delivery as a potential strategy to improve podocyte-targeted therapies in CKD.

## 1. Introduction

Chronic kidney disease (CKD) affects more than 850 million people worldwide and represents one of the leading causes of morbidity and premature mortality globally (1). Despite advances in the management of risk factors such as hypertension and diabetes, current therapies do not address the underlying cellular mechanisms of glomerular injury and patients who progress to end-stage renal disease face a lifetime of renal replacement therapy with severely impaired quality of life (2,3). There is a pressing need for mechanism-based therapeutic strategies.

Podocytes are terminally differentiated epithelial cells that form the outermost layer of the glomerular filtration barrier. Their highly specialized architecture, characterized by interdigitating foot processes bridged by the slit diaphragm, is essential for size-selective plasma filtration (4). Because podocytes cannot be replenished after injury, their structural or numerical loss is the initiating event in more than 80% of all CKD cases (2,4). Preservation of podocyte integrity therefore, represents one of the most important therapeutic objectives in nephrology.

Among the molecular mediators of podocyte injury, microRNA-21 (miR-21) occupies a central position. It is consistently upregulated in urinary exosomes of CKD patients (5) and across experimental models of glomerular injury (6). miR-21 suppresses a network of protective target genes, including the phosphatase and tensin homolog (PTEN), thereby promoting TGF-β-driven fibrosis, podocyte apoptosis and mitochondrial dysfunction (6). Therapeutic inhibition of miR-21 has shown protective effects in preclinical models of Alport syndrome, IgA nephropathy and diabetic nephropathy (7–9), establishing it as a compelling drug target.

These findings motivated the clinical development of an anti-miR-21 locked nucleic acid (LNA-21) oligonucleotide evaluated in a clinical trial in patients with Alport syndrome (10). The trial was terminated at interim analysis for futility with study design considerations primarily discussed as contributing factors. However, the underlying biological mechanistic basis for the limited clinical efficacy has not been fully established. LNA-based oligonucleotides offer high binding affinity, nuclease resistance and superior potency compared to unmodified oligonucleotides (8,11,12), making them ideally suited for miRNA inhibition in podocytes in principle. Cellular uptake, however, is one of several factors that determine the efficacy of oligonucleotide therapeutics, and delivery to post-mitotic cells such as podocytes remains challenging (10,13,14). As terminally differentiated, post-mitotic cells, podocytes are among the cell types in which internalization of free nucleic acids is particularly inefficient in the absence of a delivery (15).

Exosomes offer a biologically compelling solution to this delivery problem. These endosome-derived extracellular vesicles (30–150 nm) exploit endogenous cellular uptake pathways, protect their cargo from nuclease degradation, are low-immunogenic and can be loaded with therapeutic nucleic acids *ex vivo* (16). We recently demonstrated that directly transfected podocyte-derived exosomes efficiently deliver miRNA and siRNA cargo to cultured podocytes, achieving uptake efficiencies of up to 96.8% and producing functional target gene suppression (17). Whether this platform extends to LNA chemistry, which differs fundamentally from siRNA and miRNA in its binding mechanism and nuclease resistance, and whether it maintains efficacy under conditions of active injury, remains an open question.

Here, we systematically address these questions. We loaded podocyte-derived exosomes with a fluorescently labeled LNA anti-miR-21 (Exo-LNA-21) and evaluated delivery efficiency, functional miR-21 knockdown and downstream PTEN protein restoration across three complementary model systems: immortalized SVI podocytes, primary murine podocytes and puromycin aminonucleoside (PAN)-injured primary glomeruli from nephrin:cyan fluorescent protein (CFP) transgenic mice. Scramble-LNA controls confirmed sequence specificity throughout.

## 2. Materials and Methods

### 2.1 Cell Culture

Conditionally immortalized murine podocytes (SVI podocytes; NIPOKA GmbH, Greifswald, Germany) were used (18) and handled as previously described (17). Podocytes were maintained in RPMI-1640 (Sigma-Aldrich, St. Louis, MO, USA) supplemented with 10% FBS (Boehringer Mannheim, Mannheim, Germany), 100 U/mL penicillin and 0.1 mg/mL streptomycin (Thermo Fisher Scientific, Waltham, MA, USA). For expansion, podocytes were cultured at 33 °C and 5% CO₂ and for differentiation at 38 °C and 5% CO₂ for at least two weeks. Prior to exosome isolation, cells were washed three times with phosphate-buffered saline (PBS, Sigma-Aldrich) and medium was replaced with RPMI-1640 containing 10% exosome-depleted FBS (System Biosciences, Palo Alto, CA, USA), 100 U/mL penicillin and 0.1 mg/mL streptomycin.

### 2.2 Exosome Isolation

SVI cells were seeded at 5×10^5^ total cell number in a 75 cm^2^ culture flask and cultured in 10 mL exosome-depleted medium for 3 days. Conditioned medium was centrifuged twice at 3,000×g for 15 min at room temperature (RT). ExoQuick TC (System Biosciences; 2 mL) was added, inverted and incubated overnight at 4 °C. Exosomes were pelleted at 10,000×g for 1 h at 4 °C and resuspended in RPMI-1640 without supplements, as previously described (17).

### 2.3 Exosome Loading

Exosome transfection was performed with the Exo-Fect™ siRNA/miRNA Transfection Kit (System Biosciences) following the manufacturer’s instructions with modifications as previously described (17). mirCURY LNA™ miR-21 Inhibitor (LNA-21; 5’fluorescein amidite (FAM)-labeled; #YI04100689-ADB; QIAGEN, Venlo, Netherlands), Cy3-labeled anti-miR negative control 1 (Anti-miR-Ctrl.; #AM17011; Thermo Fisher Scientific) and scrambled LNA NC (LNA-Scramble; #YI00199006-ACA; QIAGEN) were used at 545 nM in 110 μL reactions (15 min RT, then 1 h 37 °C with 100 µL exosomes), followed by column cleanup. The free inhibitors (LNA-21 only and Anti-miR-Ctrl. only) applied gymnotically without transfection reagent and exosomes without miRNA inhibitor transfection were used as controls.

### 2.4 Exosome treatment

Prior to exosome treatment, differentiated podocytes were washed twice with 10 mL of PBS and the cells were dissociated with 3 mL of trypsin–EDTA (0.05%, 0.02%, Thermo Fisher Scientific). The trypsin–EDTA reaction was blocked by the addition of 9 mL of RPMI 1640 supplemented with exosome-depleted FBS, 100 U/mL penicillin and 0.1 mg/mL streptomycin. The cells were subsequently pelleted via centrifugation at RT and 750×g for 5 min. The pellet was subsequently eluted in 1 mL of RPMI 1640 supplemented with exosome-depleted FBS, 100 U/mL penicillin and 0.1 mg/mL streptomycin. After cell counting, 7×10^4^ cells per well were seeded in each well of a collagen IV-coated (Thermo Fisher Scientific) 6-well plate in 2 mL of RPMI 1640 supplemented with exosome-depleted FBS, 100 U/mL penicillin and 0.1 mg/mL streptomycin supplied with 2 glass coverslips each. The cells were subjected to exosome treatments after 3 days. The cleaned-up exosome elution from the transfection experiments mentioned before was applied to the corresponding wells. The cells were treated for 48 h for uptake experiments and for different durations (2 h to 48 h) to characterize uptake dynamics.

### 2.5 Transmission Electron Microscopy (TEM)

TEM was performed as previously described (17), with two modifications from the original protocol: all images were acquired using a wide-angle dualspeed CCD camera Sharpeye (Tröndle, Moorenweis, Germany) operated by ImageSP software and all micrographs were processed using Affinity Photo 2 (Serif (Europe) Ltd, Nottingham, UK). Briefly, isolated exosomes were fixed with 2% paraformaldehyde in 0.1 M sodium phosphate buffer (pH 7.5) and allowed to adsorbed onto a glow-discharged, carbon-coated holey Pioloform film on 400-mesh copper grids (Plano GmbH, Wetzlar, Germany) for 20 min on ice. Grids were transferred onto four droplets of deionized water on ice for 2 min each and finally onto a drop of staining mixture (19)for 10 min on ice. After blotting with filter paper, grids were air-dried and examined with a LEO 906 (Carl Zeiss Microscopy Deutschland GmbH, Oberkochen, Germany) at 80 kV.

### 2.6 Dynamic Light Scattering (DLS)

DLS measurements were carried out on a Zetasizer Ultra (Malvern Instruments, Herrenberg, Germany) using Zetasizer Ultra software. Samples were used either undiluted for intensity-weighted distributions (Z-average) or diluted in PBS (100 μL sample + 600 μL PBS) for number-weighted distributions and loaded into disposable macro cuvettes (10 mm path length; Brand, Wertheim, Germany). After 2 min equilibration at 25 °C, measurements were recorded at 173° backscattering (at least 12 runs per sample, ∼2 min total). Particle size distributions were determined using refractive index 1.36 and absorption coefficient 0.001 (PBS solvent settings). Both intensity-weighted and number-weighted distributions were recorded; the number-weighted distribution was used as primary readout of true particle size.

### 2.7 Confocal Laser-Scanning Microscopy (cLSM)

Confocal laser-scanning microscopy was performed on an Evident FV3000 system (Evident, Tokyo, Japan) equipped with 20× air-, 40× oil and 60× water/oil objectives. For immunofluorescence staining of PTEN and CD9, cells were washed twice with PBS and fixed with 2% paraformaldehyde (PFA) for 10 min, permeabilized with 0.3% Triton X-100 for 3 min and blocked in PBS containing 2% fetal bovine serum (FBS), 2% bovine serum albumin (BSA) and 0.2% fish gelatin for 1 h. Cells were then incubated with a rabbit anti-PTEN primary antibody (Sigma-Aldrich; Cat. No: 9552S, 1:100 dilution, 1 h RT) or rat anti-CD9 primary antibody (BD Biosciences; Cat. No: 553758, 1:100 dilution, 1 h RT), followed by incubation with Alexa Flour® 647 anti-rabbit secondary antibody (Thermo Fisher Scientific; Cat. No: A-31573, 1:300 dilution, 1 h RT) or Alexa Flour® 647 anti-rat secondary antibody (Thermo Fisher Scientific; Cat. No: A-21247, 1:300 dilution, 1 h RT). F-actin was visualized using Alexa Fluor™ Plus 488 or 555 Phalloidin (Thermo Fisher Scientific; Cat. No: A12379 or A34055, 1:100 dilution, 1 h RT) and nuclei were counterstained with DAPI (Sigma-Aldrich; 1:100 dilution, 2 min RT). Finally, samples were mounted in Mowiol (Carl Roth, Karlsruhe, Germany) prior to imaging.

All representative maximum intensity projection (MIP) images were taken using a 63× oil immersion objective and generated from Z-stacks acquired over an area of 2048 µm × 2048 µm. A 0.5 µm step size was used between optical slices, resulting in an approximate total Z-depth of 5 µm. For the quantification of mean PTEN and LNA-21 fluorescence intensities (MFI), expressed in arbitrary units per µm² (a.u./µm²), we calculated the MFIs as the integrated fluorescence intensity normalized by the cell area using ImageJ 1.54f (n = 30 cells per condition from three independent experiments).

### 2.8 Fluorescence-activated cell sorting (FACS) and Imaging Flow Cytometry

For FACS, SVI podocytes were treated for 48 h, washed twice with PBS, detached with trypsin–EDTA (0.05%, 0.02%; Thermo Fisher Scientific), pelleted at 750×g for 5 min at RT, fixed in 2% PFA for 15 min, washed twice with PBS and resuspended in 500 μL PBS as previously described (17). FACS was performed on an Aria III (BD Biosciences, Franklin Lakes, NJ, USA) equipped with a 488 nm laser (fluorescein isothiocyanate (FITC) channel). 10,000 cells were analyzed per condition and biological replicate. Imaging flow cytometry was performed on an Amnis ImageStreamX MK I (Cytek Biosciences, Fremont, CA, USA) with a 488 nm laser at 40× magnification. 1,000 cells were analyzed per condition and biological replicate.

### 2.9 Glomerulus Isolation

For the isolation of glomeruli we used transgenic nephrin:CFP mice, in which cyan fluorescent protein (CFP) is expressed specifically in podocytes under the control of a nephrin promoter fragment, as previously described (20,21). Animals were housed at 21 °C, 60% humidity, 12:12 h light–dark cycle with ad libitum access to food and water. Experiments were performed on 6-month-old mice. All animal work was conducted in compliance with national animal welfare legislation with prior approval from the competent local authority. Glomerulus isolation was performed as previously described (22) with some modifications. After collection with a 70 μm cell strainer (BD Biosciences) the isolated glomeruli were recovered by reverse-direction rinsing of the cell strainer into a Petri dish before being assessed under the microscope to ensure purity. If sufficient purity was confirmed, the contents of the Petri dish were transferred into a 15 mL conical tube and centrifuged at 300×g for 5 minutes at 20 °C. The supernatant was removed and the pellet was resuspended in 500 μL RPMI 1640 medium (Sigma‒Aldrich) supplemented with 10% fetal bovine serum (FBS; Boehringer Mannheim). Retained glomeruli were resuspended in exosome-depleted RPMI-1640 without phenol red supplemented with 10% exo-depleted FBS, 100 U/mL penicillin and 0.1 mg/mL streptomycin, seeded in uncoated 6-well plates for PAN injury experiments and in uncoated 15-well plates for live-cell imaging and cultured at 37 °C and 5% CO₂. Exosome treatment for live-cell imaging was initiated at day 0. PAN injury (12.5 µg/mL) and exosome treatment for injury experiments were initiated simultaneously at day 0; RNA was isolated after 6 days (Fig. 7 A).

### 2.10 Isolation and Culture of Primary Podocytes

For primary podocyte isolation glomeruli were cultured in 25 cm^2^ collagen IV-coated culture flasks for two weeks at 38 °C and 5% CO2 in RPMI 1640 medium (Sigma‒ Aldrich) supplemented with 10% FBS (Boehringer Mannheim), 100 U/mL penicillin and 0.1 mg/mL streptomycin (Thermo Fisher Scientific). Over this time-period primary podocytes were allowed to grow out and a progressive loss of the CFP fluorescence signal could be observed in grown out cells. After two weeks the primary cells were passaged to establish primary podocyte cultures in 25 cm^2^ collagen IV-coated flasks at 38 °C and 5% CO2 in RPMI 1640 medium (Sigma‒Aldrich) supplemented with 10% FBS (Boehringer Mannheim), 100 U/mL penicillin and 0.1 mg/mL streptomycin (Thermo Fisher Scientific). After 7 days no CFP fluorescence was detectable and the cells were dissociated with 3 mL trypsin–EDTA (0.05%, 0.02%, Thermo Fisher Scientific). The trypsin–EDTA reaction was blocked by the addition of 5 mL of RPMI 1640 supplemented with exosome-depleted FBS, 100 U/mL penicillin and 0.1 mg/mL streptomycin. The cells were subsequently pelleted via centrifugation at RT and 1000×g for 5 min. The pellet was subsequently eluted in 1 mL of RPMI 1640 supplemented with exosome-depleted FBS, 100 U/mL penicillin and 0.1 mg/mL streptomycin. After cell counting, 3×10^4^ cells per well were seeded in each well of a collagen IV-coated (Thermo Fisher Scientific) 6-well plate in 2 mL of RPMI 1640 supplemented with exosome-depleted FBS, 100 U/mL penicillin and 0.1 mg/mL streptomycin. The cells were subjected to exosome treatments after 3 days. The cleaned-up exosome elution from the transfection experiments were applied to the corresponding wells. The cells were treated for 48 h for uptake experiments (Fig. 6 A).

### 2.11 Western Blot

For exosomal protein detection, 100µL of isolated exosomes from SVI podocytes that were transfected with Anti-miR-Ctrl., LNA-21 and exosomes only were supplemented with 20 µL of ExoQuick TC, inverted and incubated on ice for 1 h. Afterwards, the samples were centrifuged for 1 h at 14,000×g and 4 °C. The pellets were eluted in Pierce RIPA Buffer (Thermo Fisher Scientific) and Halt™ Protease-Inhibitor-Cocktail (100x) (Thermo Fisher Scientific). Further protein isolation and concentration determination via the Bradford assay were performed as previously described (16,21). The exosomal protein lysates were subsequently adjusted to 20 μg/lane (TSG101) and 10 μg/lane (CD9).

Protein detection from cells was carried out as previously described (17,21). The cells that were transfected with LNA-21 only, LNA-21 with exosomes (Exo-LNA-21) and exosome only (Exo-only) were washed three times with 2 mL of PBS prior to protein isolation after 48 h. The protein lysates were adjusted to 20 μg/lane (PTEN) and 10 μg/lane (GAPDH).

All samples were mixed with 6× sample buffer (0.35 M Tris [pH 6.8], 0.35 M SDS, 30% v/v glycerol, 0.175 mM bromophenol blue) and boiled at 95 °C for 5 min. The protein samples were separated on a 4–20% gradient Mini-Protean TGX gel stain-free (Bio-Rad, Hercules, CA, USA). The separated proteins were blotted on nitrocellulose membranes via the Trans-Blot Turbo RTA Transfer Kit (Bio-Rad) and the Trans-Blot Turbo Transfer System (Bio-Rad) at 2.5 A/25 V for 5 min. Membranes were washed in 1× TBS + T wash buffer (50 mM Tris, 150 mM NaCl, 10 mM CaCl2 and 1 mM MgCl2 supplemented with 0.1% Tween-20; AppliChem, Darmstadt, Germany) and blocked in wash buffer supplemented with 5% milk powder (blocking solution) for 1 h at room temperature. The primary antibodies were diluted in blocking solution and incubated with the membranes overnight. After being washed 3 × 5 min with wash buffer, the membranes were incubated with secondary antibodies for 45 min, washed again for 4 × 5 min with wash buffer, developed with the ECL Prime Western Blotting Detection Reagent (Cytiva Europe GmbH, Freiburg, Germany) and visualized on X-ray films (Cytiva Europe GmbH) by using Carestream Kodak autoradiography GBX developer/fixer solutions. For normalization and usage of alternative antibodies on the same blot, the blots were stripped.

The following primary antibodies were used at the final concentrations: anti-TSG101 (Sigma‒Aldrich; Cat. No: T5826, 1:1000), anti-CD9 (BD Biosciences; Cat. No: 553758, 1:2000), anti-PTEN (Sigma‒Aldrich; Cat. No: 9552S, 1:1000), anti-GAPDH (Proteintech Rosemont, USA; Cat. No: 10494-1-AP, 1:10000) and secondary antibodies: anti-rabbit HRP (Santa Cruz Biotechnology, Dallas, TX, USA; Cat. No: 31463) for TSG101 (1:2000); PTEN (1:5000); GAPDH (1:20000), anti-rat HRP (Santa Cruz Biotechnology; Cat. No: 31470) for CD9 (1:10000).

### 2.12 RNA isolation and TaqMan miRNA RT-qPCR

Total RNA from immortalized podocytes, primary podocytes and isolated glomeruli was extracted using TRI Reagent (Sigma-Aldrich) according to the manufacturer’s instructions after washing twice with PBS. cDNA synthesis was performed with 10 ng of total RNA as previously described with the following kit and assays: TaqMan™ miRNA RT Kit (Thermo Fisher Scientific) and Taqman™ miRNA assays hsa-miR-21-5p (#000397; Thermo Fisher Scientific) and U6 snRNA (#001973; Thermo Fisher Scientific). The RT-qPCR experiments were performed as previously described on a QuantStudio 3 qPCR cycler (Thermo Fisher Scientific). We used the TaqMan™ Universal Master Mix II without UNG and the previously described Taqman™ assays. We conducted the following cycle scheme: 95 °C 10 min; 45 cycles of 95 °C for 15 s and 60 °C for 60 s. All samples were measured in technical triplicates in addition to biological replicates. Ct-values ≥ 38 were excluded from analysis. Fold changes were determined by the 2−ΔΔCt method and were normalized to U6 and the corresponding controls (indicated individually).

### 2.13 Statistical Analysis

Statistical analysis was performed using GraphPad Prism 11 (GraphPad Software, San Diego, CA, USA). All data were checked for Gaussian distribution using the Kolmogorov–Smirnov test. Where applicable, log₁₀ transformation was applied prior to parametric testing, as 2−ΔΔCt values are log-normally distributed. Group comparisons were performed using one-sample t-test (vs. a hypothetical mean), unpaired t-test (two groups), one-way or two-way ANOVA with Bonferroni or Tukey post hoc correction, or Brown-Forsythe ANOVA with Dunnett’s T3 post hoc test for groups with unequal variances. Non-normally distributed data were analyzed using Kruskal–Wallis with Dunn’s post hoc test or Mann–Whitney U test. All values are displayed as mean ± standard deviations (SD) with individual data points superimposed. p ≤ 0.05 was considered statistically significant.

## 3. Results

### 3.1 Exosome integrity is maintained following LNA-miR-21 transfection

To investigate the influence of the transfection on the integrity of native exosomes, we characterized the exosomes of immortalized podocyte before (Exo-only) and after direct transfections with LNA-21 (Exo-LNA-21) and with Anti-miR-Ctrl. (Exo-Anti-miR-Ctrl.). TEM analysis revealed a preserved cup-shaped morphology which is typical for exosomes and consistent particle sizes of approximately 20-50 nm in all three treatment groups (Fig. 1A). Additionally, we observed intact and homogenous exosomes throughout the different groups. Compared to both other groups, Exo-only showed a slightly higher degree of particle aggregation.

**Figure 1.**
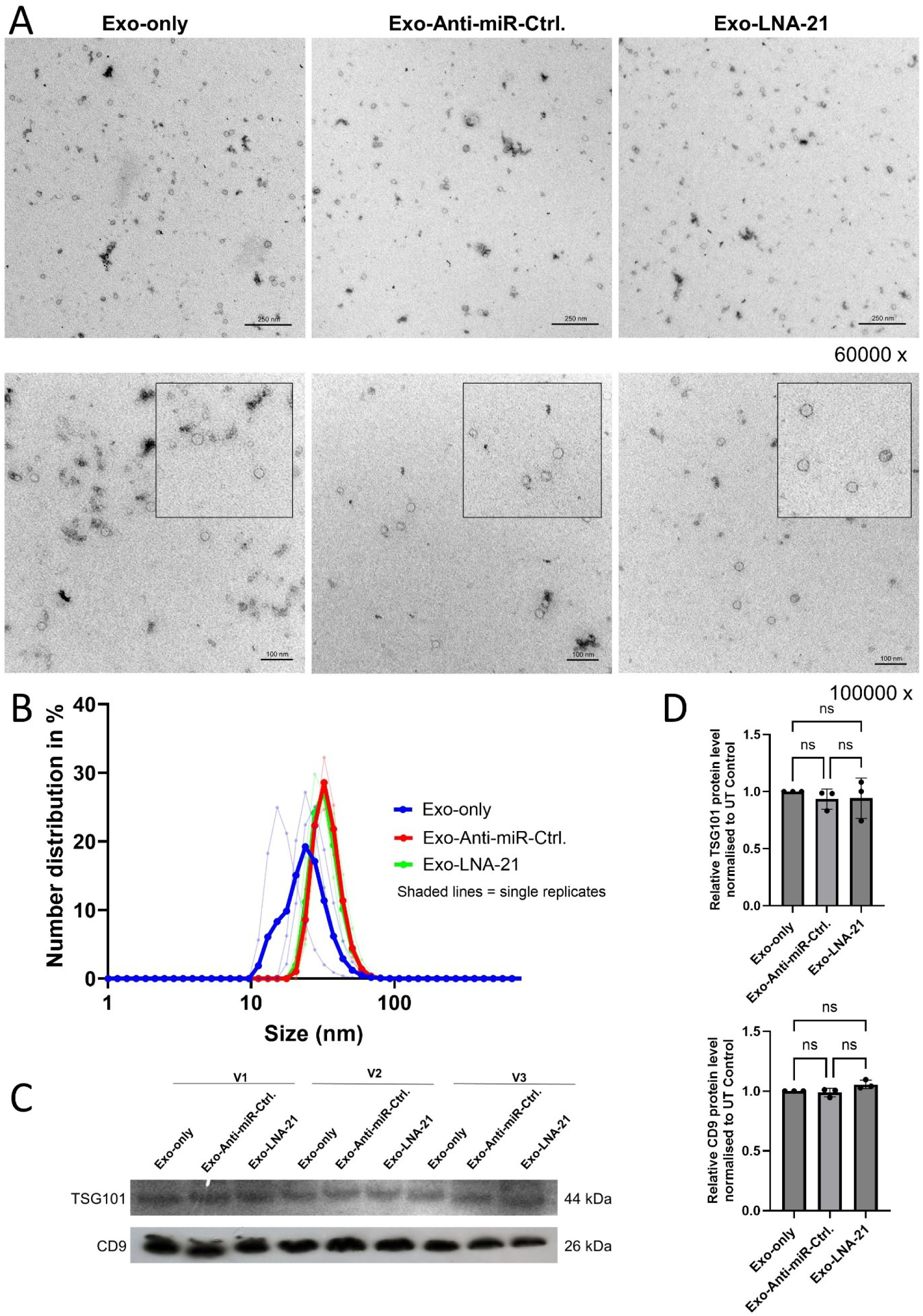
Exosome characterization before and after transfection. (A) TEM micrographs of Exo-only, Exo-Anti-miR-Ctrl. and Exo-LNA-21 at 60,000× magnification (upper) and 100,000× magnification (lower). Scale bars = 250 nm (upper) and 100 nm (lower). (B) DLS number-weighted size distributions for Exo-only (blue), Exo-Anti-miR-Ctrl. (red) and Exo-LNA-21 (green). Thick lines: means (n = 3); shaded lines: individual preparations. (C) Western blot for TSG101 (44 kDa) and CD9 (26 kDa) in Exo-only, Exo-Anti-miR-Ctrl. and Exo-LNA-21 across three independent preparations (V1–V3). (D) Quantification of TSG101 and CD9 protein levels normalized to Exo-only control. No significant differences were visible between the treatment groups. Bar graphs: mean ± SD; dots: n = 3. All comparisons n.s. (One-way ANOVA).

To validate these results, we performed DLS measurement. The number-weighted size distribution revealed comparable dominant peaks at 20–35 nm for Exo-only, Exo-Anti-miR-Ctrl. and for Exo-LNA-21 (Fig. 1B), confirming the observed exosome size obtained by TEM. Additionally, these observations corroborate that the exosomal population itself remains unchanged after transfection. To confirm that LNA loading did not compromise exosomal marker expression, Western blot analysis for TSG101 and CD9 was performed. Thereby we have not identified significant differences across all three conditions (Fig. 1C–D). Together, these data demonstrate that the loading of exosomes with LNAs maintains exosome number, integrity and marker profile.

### 3.2 Exosome-delivered cargo is taken up by SVI podocytes in a time-dependent manner

To determine the relationship between LNA–miR-21-loaded exosomes and the duration of uptake and processing, we analyzed the time-dependent internalization of exosome-delivered cargo in SVI podocytes by using FAM-labeled Exo-LNA-21 (Fig. 2 A, green) or Cy3-labeled Exo-Anti-miR-Ctrl. (Fig. 2 B, magenta). The treated cells were imaged by cLSM after 2 h, 4 h, 24 h and 48 h. The intracellular fluorescent signals were detectable as early as 2 h and increased progressively, with pronounced perinuclear accumulation at 24 h and 48 h in both conditions (Fig. 2 A and B).

**Figure 2.**
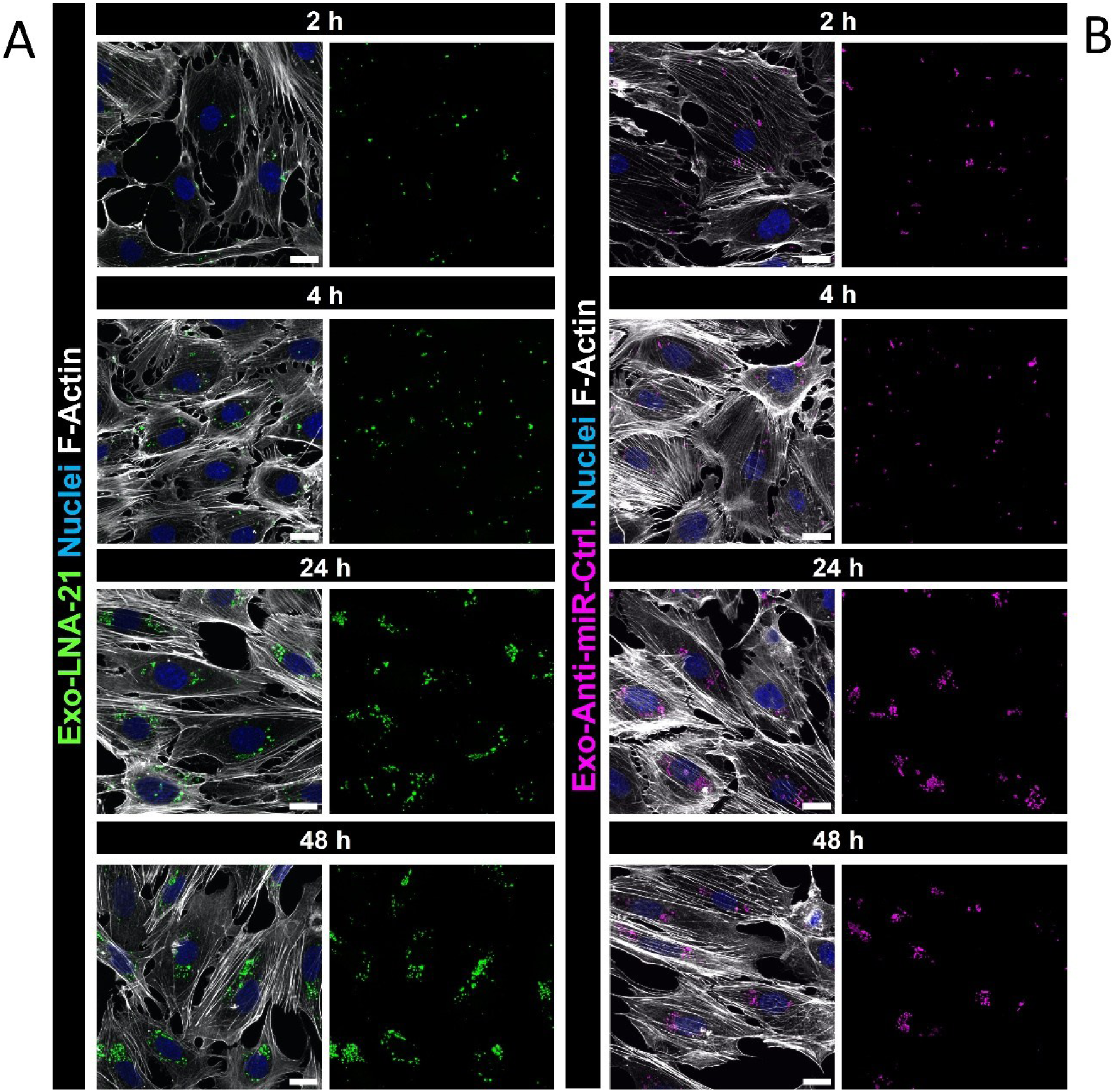
Time-course of exosomal cargo uptake in SVI podocytes by cLSM. (A) Exo-LNA-21 (green: 5’FAM-LNA; grays: F-Actin; blue: Nuclei) at 2 h, 4 h, 24 h and 48 h. (B) Exo-Anti-miR-Ctrl. (magenta: Cy3-anti-miR; grays: F-Actin; blue: Nuclei) at the same time points. Intracellular fluorescent signal was detectable as early as 2 h and increased progressively over 48 h, with strong perinuclear accumulation at 24 and 48 h in both Exo-LNA-21- and Exo-Anti-miR-Ctrl.-treated cells. Scale bars = 20 μm.

### 3.3 Exosome-mediated LNA-21 delivery induces efficient uptake and near-complete miR-21 knockdown in cultured podocytes

To compare free versus exosome-mediated LNA-21 and anti-miR-Ctrl. delivery into immortalized SVI podocytes, we first examined the uptake by confocal immunofluorescence 48 h after transfection (Fig. 3 A). In Exo-Anti-miR-Ctrl.- and Exo-LNA-21-treated cells, the cargo signals were clearly visible in the perinuclear cytoplasm as described above. By contrast, free applied Anti-miR-Ctrl. only and LNA-21-only as controls showed no detectable intracellular fluorescence signals, demonstrating that cellular uptake is strictly dependent on exosomal delivery. To quantify these results, we performed FACS analysis 48 h after treatment. Exo-LNA-21-transfected podocytes showed 76.6% FAM-positive cells compared to only 0.82% in LNA-21-only, 0.03% in Exo-only and 0.13% in untreated cells (Fig. 3 C). Imaging flow cytometry confirmed that the FAM fluorescence signal was localized intracellularly rather than being surface-bound (Fig. 3 D). Cytoplasmic LNA accumulation was detected exclusively in Exo-LNA-21-treated cells, whereas cells treated with free LNA-21, exosomes alone, or left untreated showed no intracellular signal (Fig. 3 D). Consistent with these results, imaging flow cytometry showed 79.75% of podocytes were FAM-positive following Exo-LNA-21 treatment, compared with 0.05% in LNA-only, 0.03% in Exo-only and 0.00% in untreated cells.

**Figure 3.**
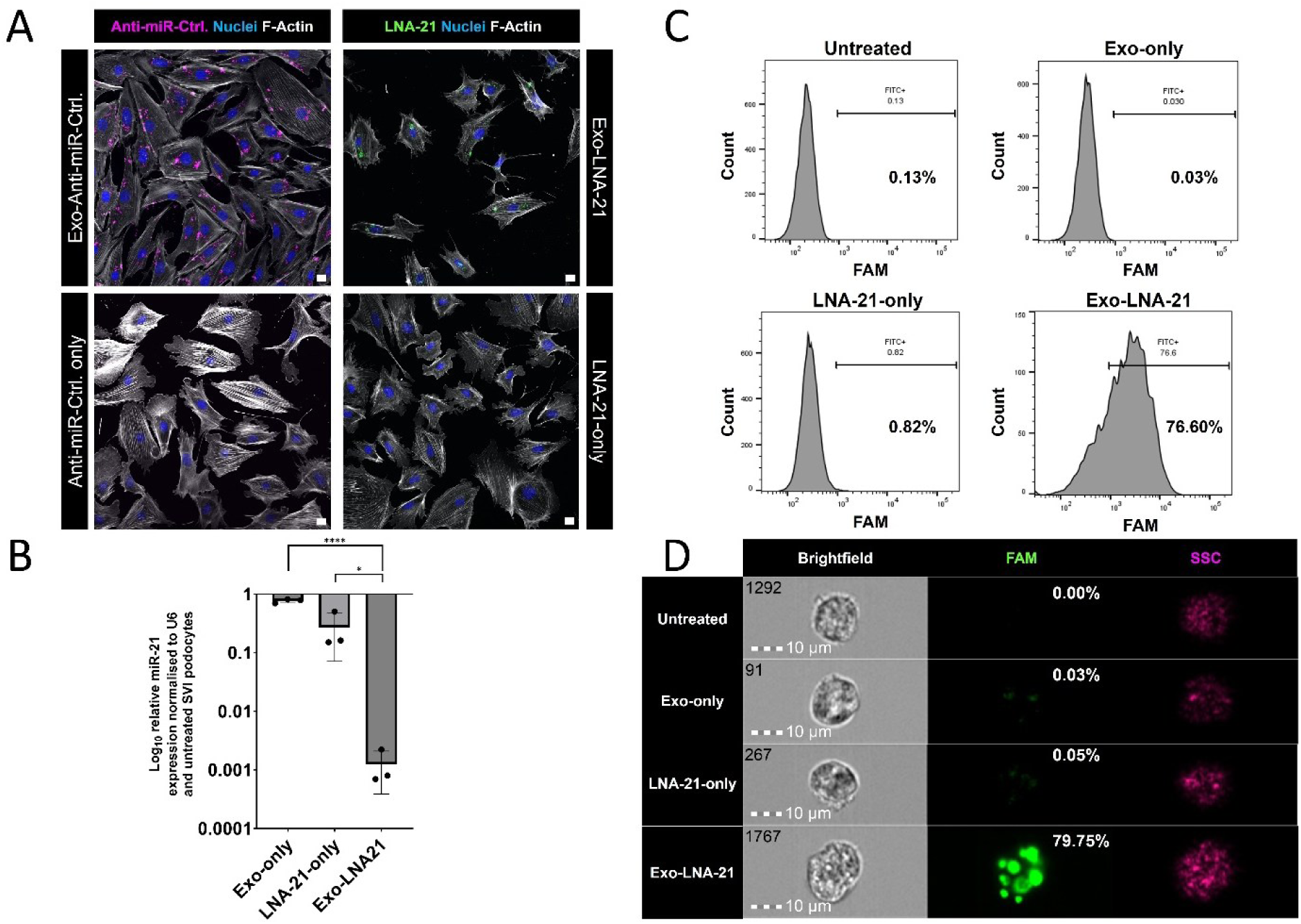
Exosome-mediated LNA–miR-21 uptake induces functional miR-21 knockdown in SVI podocytes. (A) Immunofluorescence of Exo-Anti-miR-Ctrl. (upper left) and Exo-LNA-21 (upper right) vs. Anti-miR-Ctrl. only (lower left) and LNA-21-only (lower right). Grays: F-Actin; green: FAM-labeled LNA-21(right); magenta: Cy3-labeled Anti-miR-Ctrl.(left); blue: Nuclei. Perinuclear cargo signals were detected exclusively in treated Exo-Anti-miR-Ctrl.- and Exo-LNA-21-treated cells, confirming that efficient uptake requires exosomal delivery. Scale bars = 10 μm. (B) miR-21 expression (2−ΔΔCt, log₁₀-scale) in SVI podocytes (48 h, normalized to U6 and untreated). Bar graph: mean ± SD; dots: n = 3. TaqMan RT-qPCR showed an 804-fold miR-21 reduction with Exo-LNA-21 versus Exo-only and 3.5-fold with free LNA-21-only (**** p < 0.0001 and * p = 0.046, respectively), demonstrating a > 200-fold higher delivery-dependent difference in knockdown efficiency. Statistical analysis Welch’s t-test. (C) FACS histograms at 48 h showing FAM-positive (FITC+) cells (%) for untreated, Exo-only, LNA-21-only and Exo-LNA-21 (n = 3; 10000 cells each; merged IFC value shown). FACS analysis revealed 76.60% FAM-positive cells in Exo-LNA-21-treated podocytes, compared to 0.82% (LNA-21-only), 0.03% (Exo-only) and 0.13% (untreated). (D) Imaging flow cytometry confirmed intracellular localization of the fluorescence signal, with 79.75% FAM-positive cells in the Exo-LNA-21 group compared to 0.05% (LNA-21-only), 0.03% (Exo-only) and 0.00% (untreated; Fig. 3 D). Brightfield, FAM (green) and SSC (magenta) channels for each condition (n = 3, 1000 cells each). Scale bars = 10 μm.

To confirm functional cargo functionality, we performed RT-qPCR demonstrating that Exo-LNA-significantly downregulated miR-21 by 804-fold in cultured podocytes (p < 0.0001) compared to Exo-only (0.78-fold). In contrast, treatment with free LNA-21 alone resulted in a significantly smaller, 3.5-fold reduction (p = 0.0460; Fig. 3B). This shows a more than 200-fold difference in knockdown efficiency between exosomal and free LNA-21 delivery.

### 3.4 Exosome-mediated LNA-21 delivery restores PTEN protein expression

To prove the functional downstream activity of the delivered LNA-21 at protein level, we investigated the expression of a miR-21 target PTEN by immunofluorescence microscopy. Exo-LNA-21-treated podocytes showed significantly higher PTEN MFI compared to both LNA-21-only and Exo-only controls (p = 0.0003; Fig. 4 A and B).

**Figure 4.**
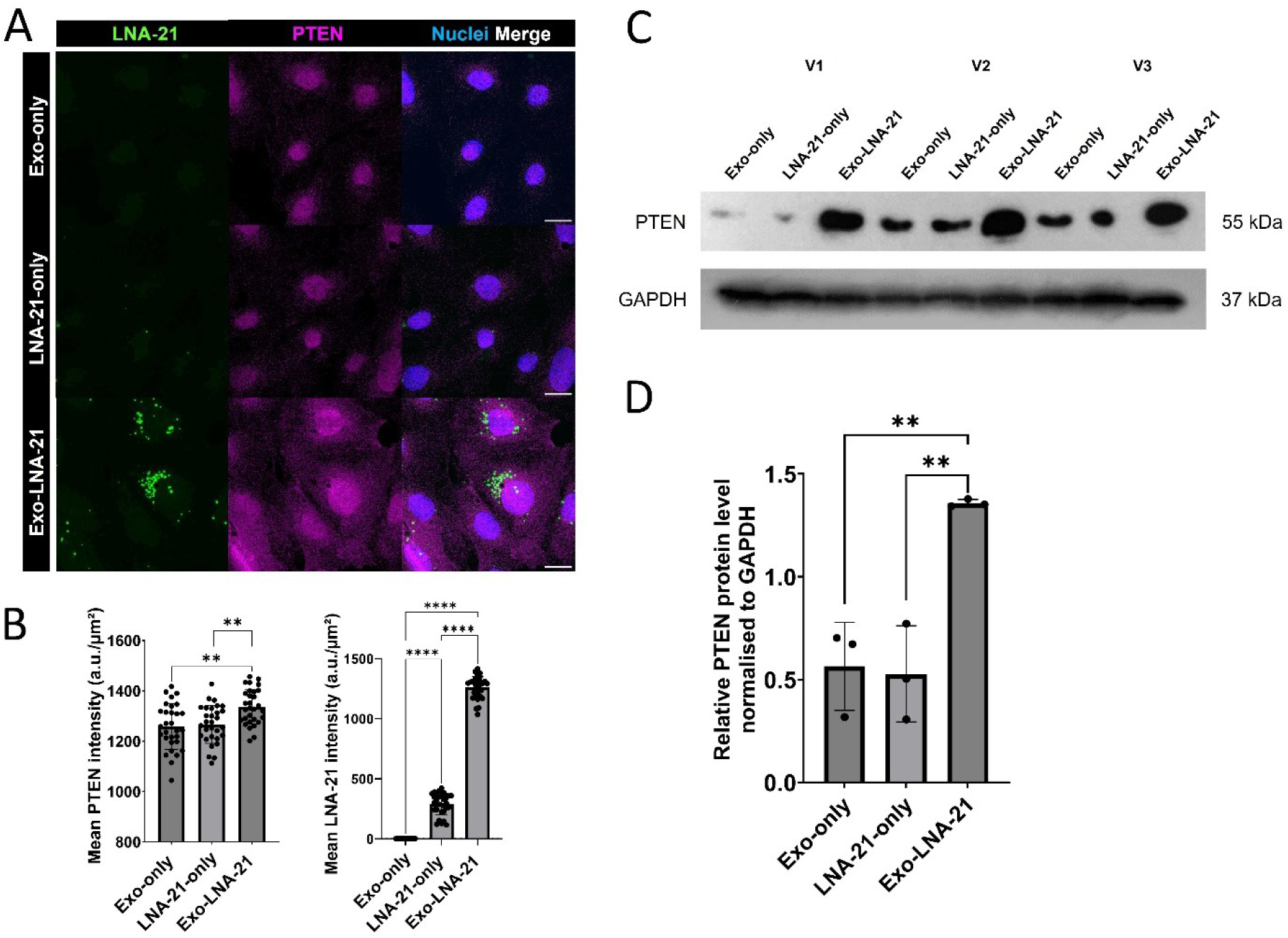
Exo-LNA-21 restores PTEN protein expression. (A) Confocal immunofluorescence of SVI podocytes treated with Exo-only, LNA-21-only, or Exo-LNA-21. Green: LNA-21; magenta: PTEN; blue: Nuclei; right column: merge. Scale bars = 10 µm. (B) Left: quantification of mean PTEN fluorescence intensity (n = 30 cells/3 experiments). ** p = 0.0013 (Exo-only vs. Exo-LNA-21), ** p = 0.0013 (LNA-21-only vs. Exo-LNA-21; Brown-Forsythe ANOVA test with Dunnett’s T3 multiple comparisons test). Right: quantification of mean LNA-21 fluorescence intensity. **** p < 0.0001 for all pairwise comparisons (Kruskal-Wallis test with Dunn’s multiple comparisons test). (C) PTEN Western blot across three independent preparations (V1–V3). PTEN: 55 kDa; GAPDH: 37 kDa. (D) Quantification of PTEN/GAPDH ratio. * p = 0.0383 (Exo-only vs. Exo-LNA-21) and * p = 0.0395 (LNA-21-only vs. Exo-LNA-21; n = 3, One-way ANOVA with Tukey’s multiple comparisons test).

CLSM images showed distinct PTEN (magenta) and LNA-21 (green) signals, with PTEN clearly higher expressed in Exo-LNA-21-treated cells. Quantification of intracellular LNA-21 fluorescence confirmed significantly higher cargo signals in Exo-LNA-21-treated cells compared to LNA-21-only and Exo-only controls (p < 0.0001, respectively; Fig. 4 A and B).

To confirm these findings, we performed Western blot analysis which revealed that Exo-LNA-21 significantly increased the PTEN expression (1.357-fold) compared to Exo-only (0.564-fold; p = 0.0383) and free LNA-21-only (0.527-fold; p = 0.0395). There was no significant difference between Exo-only and LNA-only (p = 0.850; Fig. 4 C and D).

To confirm sequence specificity, we directly compared an Exo-LNA-Scramble control with Exo-LNA-21 by immunofluorescence. Interestingly, only Exo-LNA-21 reduced PTEN levels (MFI = 1643; p = 0.0005; Fig. 5 A and B) compared to Exo-LNA-Scramble (MFI = 1603), demonstrating that the PTEN changes reflect specific miR-21 inhibition and not a non-specific exosomal effect. This was further corroborated at the RNA level by RT-qPCR, where Exo-LNA-21 led to a ∼97% reduction in miR-21 relative to Exo-LNA-Scramble (0.034-fold; p = 0.0008; Fig. 5 C).

**Figure 5.**
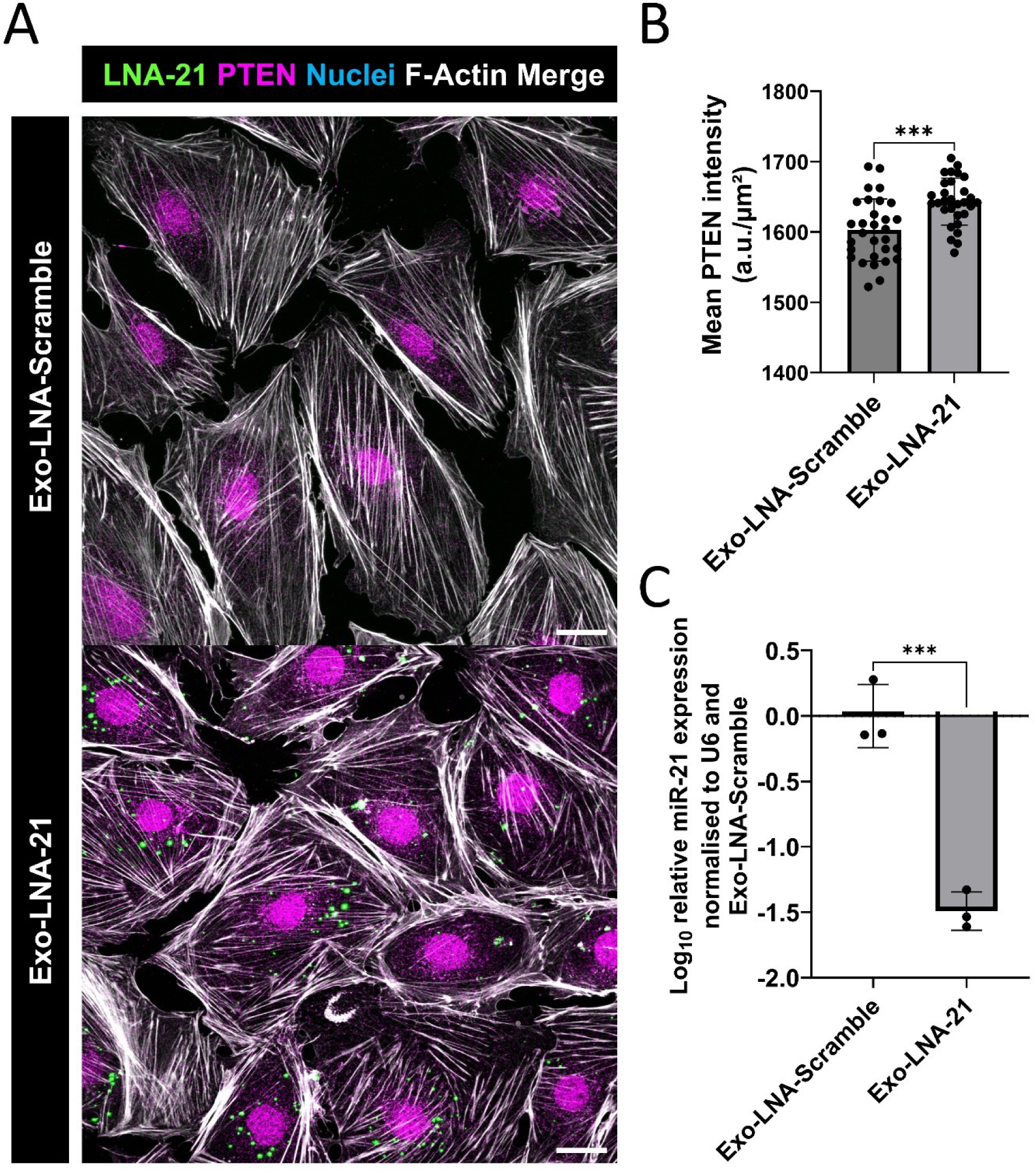
Sequence specificity of Exo-LNA-21-mediated miR-21 inhibition. (A) Confocal images of Exo-LNA-Scramble vs. Exo-LNA-21 (green: LNA-21; magenta: PTEN; white: F-Actin; blue: Nuclei). Scale bar = 10 μm. (B) PTEN MFI quantification of Exo-LNA-Scramble vs. Exo-LNA-21 (n = 30 cells/3 experiments; *** p = 0.0005, Mann-Whitney test). (C) miR-21 expression normalized to Exo-LNA-Scramble (n = 3; *** p = 0.0008; Unpaired t-test).

**Figure 6.**
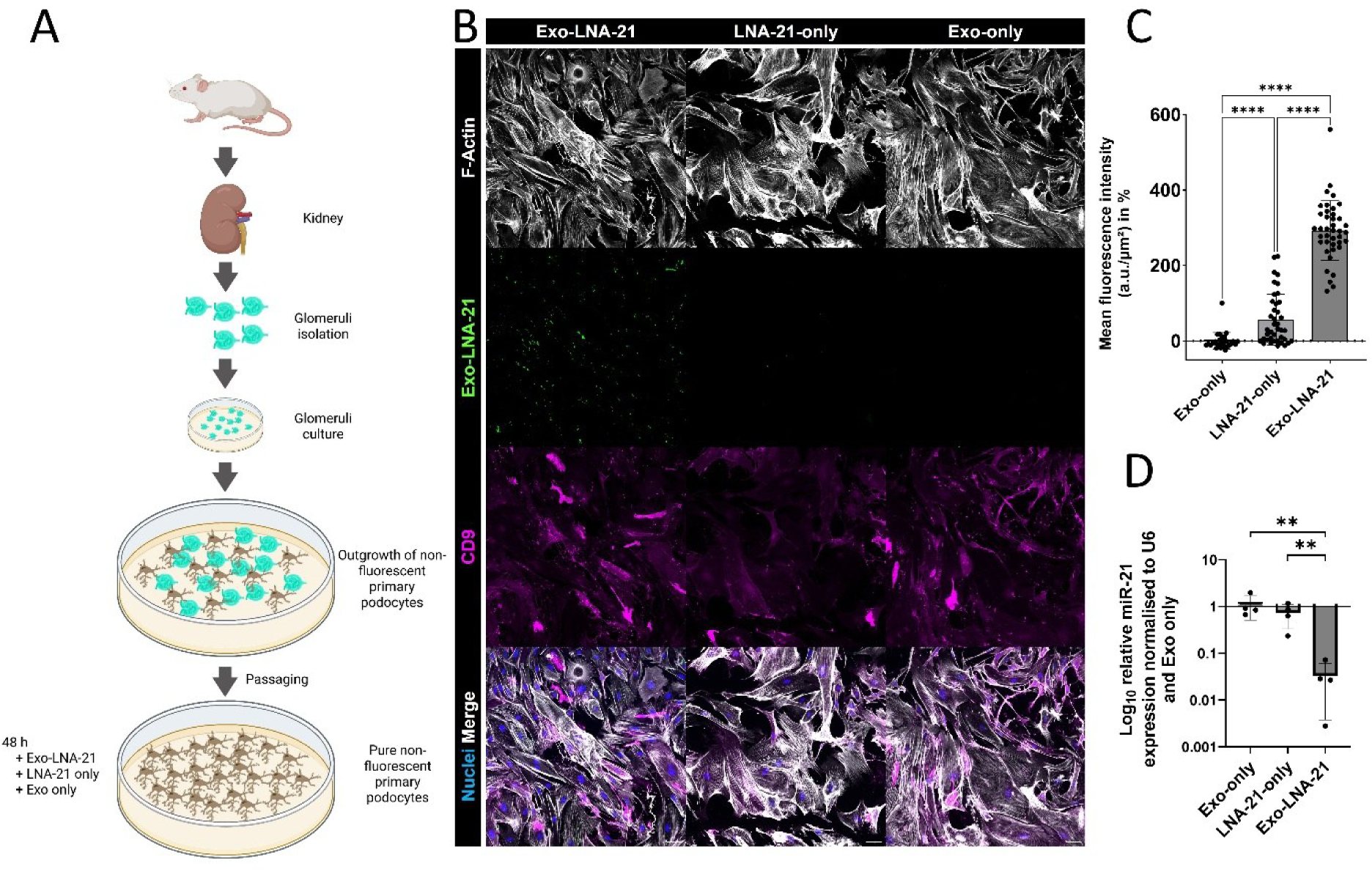
Exosomal LNA-21 delivery and miR-21 knockdown in primary murine podocytes. (A) Schematic overview of the experimental setup. Created in BioRender. Lange, T. (2026) https://BioRender.com/07zpeeb. (B) Immunofluorescence of primary murine podocytes (Exo-only, LNA-21-only, Exo-LNA-21) stained for F-Actin (white), FAM-LNA-21 (green), CD9 (magenta) and Nuclei (blue) with merged channels. Scale bar = 20 μm. (C) FAM-LNA-21 fluorescence intensity per cell area and number (a.u.); n = 30 cells from 3 independent primary podocyte preparations. **** p < 0.0001(Brown-Forsythe ANOVA test with Dunnett’s T3 multiple comparisons test). (D) miR-21 expression (2−ΔΔCt, log₁₀-scale, normalized to U6 and mean Exo-only ΔCt across V1–V4) in primary murine podocytes (48 h, n = 4 independent preparations). Mean ± SD; dots: individual preparations. Exo-LNA-21 vs. Exo-only ** p = 0.0050; Exo-LNA-21 vs. LNA-21-only * p = 0.0404 (One-way ANOVA with Uncorrected Fisher’s LSD).

**Figure 7.**
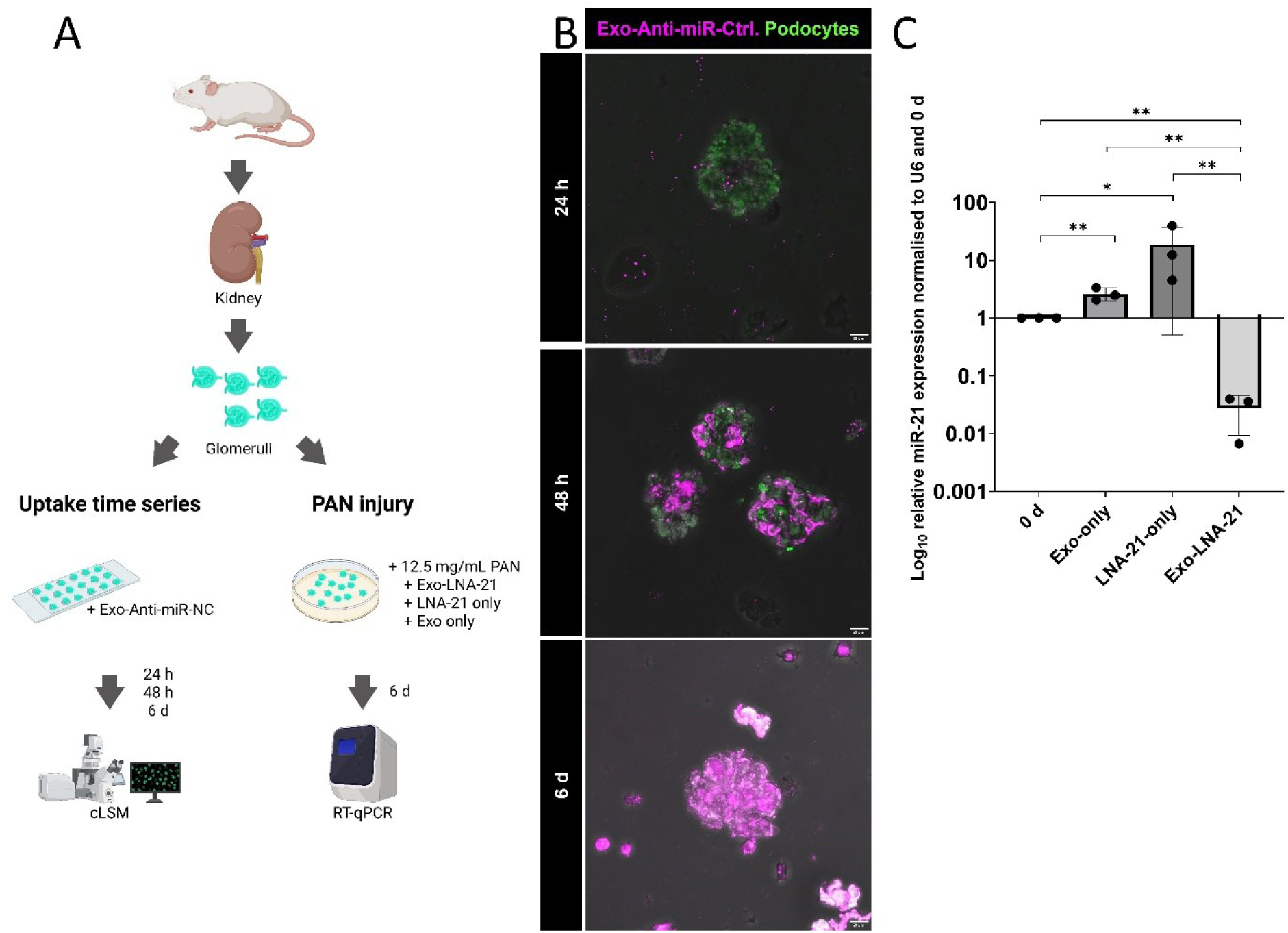
Exosomal Anti-miR-Ctrl. uptake by isolated glomeruli under healthy and PAN-injured conditions. (A) Schematic overview of the experimental design. Created in BioRender. Lange, T. (2026) https://BioRender.com/8yiqfbd. (B) cLSM time series of glomeruli treated with Exo-Anti-miR-Ctrl. (Cy3: magenta; Podocytes CFP: green; brightfield overlay) at 24 h, 48 h and 6 d. Scale bars = 20 μm. (C) miR-21 expression (2−ΔΔCt, log10-scale, normalized to U6 and Day-0) in PAN-injured glomeruli treated for 6 d. n = 3 independent experiments. ** p = 0.003 (Exo-LNA-21 vs. Day-0); ** p = 0.003 (Exo-only vs. Day-0); * p = 0.015 (LNA-21-only vs. Day-0); *** p = 0.002 (LNA-21-only vs. Exo-LNA-21).

### 3.5 LNA-21 generated a potent miR-21 knockdown in primary murine podocytes

To validate the delivery platform with primary cells that are difficult to transfect, we studied the delivery of LNA-21 in primary murine podocytes (Fig. 6 A).

Immunofluorescence staining of primary podocytes for the exosome marker CD9, for Exo-LNA-21, and F-Actin showed that LNA-21 cargo colocalized with the CD9-positive vesicles in the perinuclear region (Fig. 6 B, Suppl. Fig. 1). This is consistent with the endocytic uptake pathways we previously described for siRNA and miRNA cargo (17). Quantification of FAM-LNA-21 fluorescence intensity per cell area and number confirmed significantly higher intracellular signals in Exo-LNA-21-treated primary podocytes (MFI = 292.7%) compared to LNA-21-only (MFI = 56.44%; p < 0.0001) and Exo-only (MFI = 8.663*10^-6^%; p < 0.0001; Fig. 6 C).

To proof the functional knockdown in primary podocytes, RT-qPCR was performed across four independent preparations. Exo-LNA-21 produced a consistent and significant 0.033-fold reduction in miR-21 relative to Exo-only in all four experiments (p = 0.0050; Fig. 6 D). In contrast, free LNA-21-only showed no significant miR-21 suppression (0.721-fold; p = 0.3022). In direct comparison, the difference between Exo-LNA-21 and LNA-21-only was highly significant (p = 0.0404). These results confirm that the delivery advantage of exosomal encapsulation is not a cell line artifact but a genuine property of primary post-mitotic podocytes.

### 3.6 Exo-LNA-21 suppresses miR-21 expression in PAN-injured glomeruli

To test the delivery platform under *ex vivo* conditions with *in situ* podocytes, glomeruli from nephrin:CFP mice were isolated as described for the GlomAssay and cultured in 15-well plates (Fig. 7 A). To assess exosome uptake by podocytes *in situ* within the intact glomerular structures, the uptake of Exo-Anti-miR-Ctrl. was assessed by cLSM at 24 h, 48 h and 6 d (Fig. 7 B). Z-stack imaging was performed with the vessel bottom plane excluded from image acquisition to prevent signal contamination from surface-adhered exosomes; corresponding images including the bottom plane are provided in Suppl. Fig. 2. The intracellular fluorescent signals were detectable within the glomeruli already at 24 h after transfection and increased progressively which confirmed the uptake of the exosomal cargo by primary podocytes *in situ*.

To extend these findings to a disease-relevant injury context, glomeruli were exposed to PAN (12.5 µg/mL) simultaneously with exosome treatment at day 0. After 6 days, RT-qPCR showed that Exo-LNA-21 achieved a highly significant reduction by ∼97% of miR-21 expression compared to Day 0 (p = 0.003; Fig. 7 C). In contrast, free LNA-21-only induced miR-21 expression by 18.8-fold (p = 0.015), which was consistently observed across all three independent experiments. The difference between Exo-LNA-21 and LNA-21-only was highly significant (p = 0.002). Exosomes without a cargo (Exo-only) resulted in a significant upregulation by 2.6-fold compared to Day 0 (p = 0.003).

## 4. Discussion

In the past, exosomes have been shown to be effective carriers for the delivery of RNAs to various cell types, including podocytes (23,24). Compared to lipid nanoparticles, which have an established delivery track record, exosomes have a higher biocompatibility, lower immunogenicity and higher biodistribution (14,25). However, achieving high transfection rates and minimizing off-target effects has remained a persistent challenge. In our recent manuscript, we were able to overcome this limitation by using directly transfected podocyte-derived exosomes, which enabled the efficient delivery of fluorescently labeled siRNA and miRNA to immortalized podocytes *in vitro* (17). The present study extends this platform to LNA anti-miR technology. Building on these previous findings, the present study demonstrates that podocyte-derived exosomes can also be used to achieve efficient miRNA knockdown using LNA-modified anti-miRNA oligonucleotides. This is particularly important because LNA oligonucleotides differ fundamentally from siRNA and miRNA in their mode of action, binding kinetics, and nuclease resistance (12). Therefore, it remained unclear whether exosomal loading and delivery would preserve their pharmacological properties and biological function. The present study addresses this question across three model systems of increasing biological complexity.

In this context, we first investigated whether LNA loading affects the exosome integrity. To this end, TEM analysis was performed to evaluate whether the loading with LNA affects exosome size and morphology. Here we show that the typically cup-shaped morphology is preserved with and without loading of LNA. However, slightly more particle aggregation was observed in the Exo-only group. This is consistent with our previous finding that the additional purification step, which was included to remove residual loading reagents and clean up cargo-loaded exosomes, resulted in a more dispersed exosome population with fewer aggregates (17). Furthermore, this was confirmed by the number-weighted distribution analysis via DLS, which accurately reflects the true major particle distribution (26). It showed indistinguishable peaks at 20–35 nm across all conditions, consistent with the homogeneous populations visible by TEM. To further assess whether LNA loading affects the expression of typical exosomal marker proteins, we performed Western blot analyses for CD9 and TSG101. (16) The results demonstrated that LNA anti-miR loading did not alter the expression of these exosomal markers. In conclusion, our loading method preserved both the structural integrity and the total number of exosomes. Such post-loading validation remains essential for robust reproducibility. While this critical aspect is often debated, it is rarely verified in most studies (27).

Using complementary approaches, including confocal microscopy, FACS, imaging flow cytometry, and RT-qPCR, we further demonstrated that exosomal encapsulation enhances intracellular uptake and delivery of LNA oligonucleotides. CLSM revealed that both Exo-LNA-21 and Exo-Anti-miR-Ctrl. cargo followed the same progressive perinuclear accumulation pattern that we previously described for siRNA and miRNA (17), with signals detectable as early as 2 h and increasing at 24 h and 48 h. These results demonstrate that this exosome platform supports efficient delivery of different oligonucleotides into podocytes. In contrast, freely applied LNA showed no detectable intracellular signal at 48 h which was confirmed by FACS quantification. By imaging flow cytometry, we independently confirmed that the Exo-LNA-21 signal was intracellular rather than surface-bound, reproducing the uptake advantage at 79.75% versus 0.05% (LNA-21-only). The convergence of all three imaging methods provides a robust and artefact-independent confirmation of genuine intracellular delivery. Furthermore, RT-qPCR corroborated these findings functionally: Exo-LNA-21 achieved an 804-fold miR-21 reduction compared to a 3.5-fold reduction with free LNA-21-only. This effect was described before for antisense oligonucleotides in neuronal cells by Yang et al. (28). Notably, free LNA-21 exerted a measurable RNA effect, demonstrating that a minor fraction enters cells gymnotically. This aligns with previous findings that different cell types (29,30) including podocytes (31), take up inhibitory oligonucleotides without transfection reagents.

Beyond assessing efficient uptake and preserved exosome integrity, we next examined whether the delivered LNA oligonucleotides remained functionally active. For this purpose, we selected PTEN, a known miR-21-regulated target protein, as a functional readout. PTEN is a well-established direct target of miR-21 in podocytes and has been shown to regulate podocyte motility and cytoskeletal integrity. (8) We found a significant restoration of PTEN after Exo-LNA-21 incubation at both immunofluorescence and Western blot, whereas this effect was not observed with free LNA-21-only. Even though we observed a slight 3.5-fold miR-21 reduction in LNA-21-only treated samples, this level of delivery was insufficient to produce any detectable change in PTEN protein, while Exo-LNA-21 significantly restored PTEN protein expression. Only the specific Exo-LNA-21 rather than the scrambled control reduced PTEN expression, confirming that the observed effects are driven by sequence-specific miR-21 inhibition. This control is particularly important given that endogenous exosomal miRNA cargo can itself influence gene expression in recipient cells (32). The fact that an LNA, with its inherently high target affinity and nuclease stability (12), requires exosomal encapsulation to exert functional effects in podocytes *in vitro* further underscores that the delivery barrier, rather than the oligonucleotide chemistry, is the rate-limiting factor in this cell type. These findings highlight the potential of this exosome-based delivery platform as a strategy to modulate downstream signaling pathways that may contribute to podocyte protection in future therapeutic applications.

To validate the results obtained in immortalized mouse podocytes in a more physiologically relevant setting, we performed the same experimental approach in primary murine podocytes isolated from a transgenic mouse strain expressing cyan fluorescent protein specifically in podocytes. In these primary podocytes, we detected a co-localization of the LNA cargo with CD9-positive perinuclear vesicular structures. Moreover, the functional miR-21 knockdown of approximately 97% was fully reproduced, confirming efficient exosome-mediated LNA delivery and activity in primary podocytes.

Since primary podocytes progressively lose CFP fluorescence in culture due to silencing of the promoter driving CFP expression, we additionally assessed FAM fluorescence as a direct readout of LNA cargo uptake. The clear detection of FAM fluorescence in Exo-LNA-21-treated primary podocytes therefore confirms genuine intracellular cargo accumulation. These findings further demonstrate that the delivery advantage of Exo-LNA-21 is not a cell-line artifact, but rather represents a reproducible property of primary murine podocytes.

We next extended our analysis to podocytes *in situ*. As previously demonstrated by our group, isolated glomeruli represent a well-suited model to study podocyte behavior while preserving their attachment to the glomerular capillary tuft. To study podocytes within their native microenvironment, we again used the transgenic mouse strain with podocyte-specific cyan fluorescent protein expression and isolated glomeruli from these animals (20,21). Isolated glomeruli were treated with exosomes under the same experimental conditions used for the cell culture experiments. Under these conditions, podocytes showed pronounced uptake of exosome-associated cargo.

Since these conditions reflect a healthy state, we next sought to investigate what happens when podocytes are shifted toward a diseased phenotype by treatment with PAN. PAN-induced podocyte injury mimics key features observed in several glomerulopathies and has been reported to be associated with a marked upregulation of miR-21 (33–35). Remarkably, Exo-LNA-21 treatment of the isolated glomeruli resulted in an approximately 97% suppression of miR-21. This effect was achieved despite the disease-associated transcriptional counterforce induced by PAN, indicating that exosomal encapsulation enables efficient LNA delivery and functional miR-21 inhibition even under active injury conditions.

## 5. Conclusions

Our study establishes exosome-based delivery as an effective strategy to render LNA anti-miR-21 functionally active in podocytes both *in vitro* and *in situ*. By overcoming the limited cellular uptake of free LNA, Exo-LNA-21 enabled potent and sequence-specific miR-21 suppression, restored PTEN expression, and remained effective even in PAN-injured isolated glomeruli. Since miR-21 plays a central role in podocyte injury and glomerular disease, these findings highlight exosome-mediated LNA delivery as a promising platform for targeted miRNA inhibition and the modulation of disease-relevant pathways in podocytes. This provides a solid foundation for preclinical studies with high translational potential for therapeutic approaches targeting glomerular diseases.

## Author Contributions

N.E. contributed to supervision. N.E. and T.L. contributed to project administration. T.L. and N.E. did the conceptualization. T.L. and N.E. contributed to funding acquisition. T.L., J.E.C.S., L.K.F. and P.L.G. performed all exosome transfection experiments. Downstream experiments including RT-qPCR, cLSM, and Western blot were performed by T.L., J.E.C.S., L.K.F., P.L.G and C.W. R.S. performed transmission electron microscopy experiments. T.L., P.L.G. and D.B. performed flow cytometry experiments. DLS experiments were done by U.J. and M.D. T.L., J.E.C.S. and L.K.F. performed data analysis and figure design. The manuscript writing was performed by T.L., J.E.C.S. and L.K.F. and N.E.. All authors proofread the manuscript.

## Funding

Our project was supported by the Federal Ministry of Education and Research (BMBF, grant 01BIH_nTTP-GCT_26/03; Tim Lange) and by the Federal Ministry of Education and Research (BMBF, grant 01GM2202B, STOP-FSGS; Prof. Nicole Endlich). Additionally, funding was provided by the Federal Ministry for Economic Affairs and Climate Action (BMWi, grant 16KN077229, project title: Alterna Tier-vivoPod; Prof. Nicole Endlich). In addition to this, we received startup funding from the *“Forschungsverbünde FVCM, FVMM, GANI_MED, Digital Health Lab”* of the University Medicine Greifswald (Tim Lange). Further financial support was received from the Dr. Gerhard Büchtemann Fund, Hamburg, Germany and the Südmeyer Foundation for Kidney and Vascular Research (“Südmeyer Stiftung für Nieren- und Gefäßforschung”).

## AI Use Statement

During the preparation of this manuscript, the authors used Claude (Anthropic) and ChatGPT (OpenAI) to assist with language editing and manuscript revision. All scientific content, data, analyses and conclusions were generated and verified by the authors. The authors take full responsibility for the integrity and accuracy of the published work.

## Conflicts of Interest

N.E. is CEO of the NIPOKA GmbH, Greifswald, Germany. N.E. and T.L. filed a patent that is related to this study.

## Acknowledgement

The authors thank Annette Meuche for excellent technical assistance regarding electron microscopy.

---

**Supplementary Figure 1.**
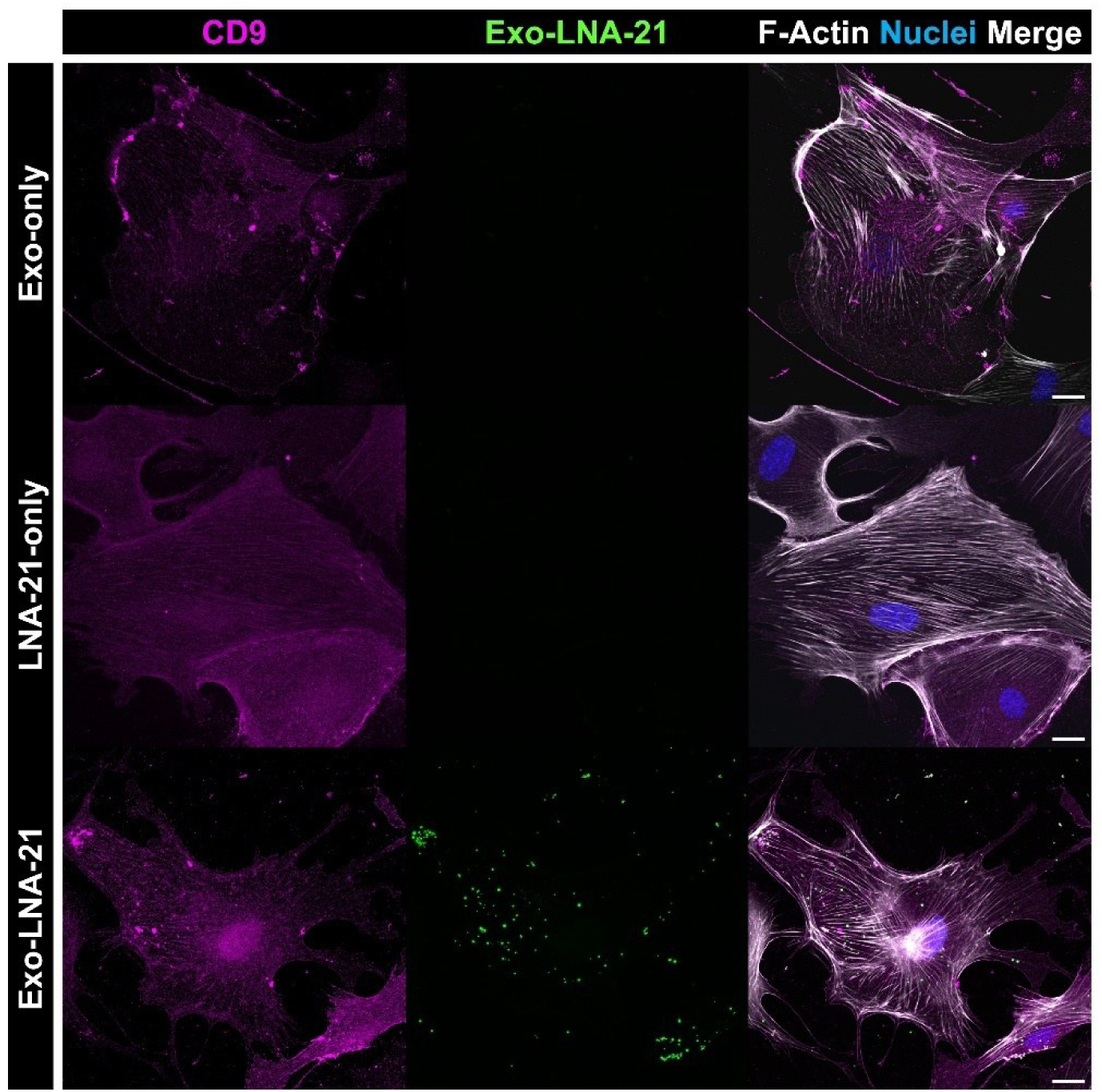
Exosomal LNA-21 delivery in primary murine podocytes. Immunofluorescence of primary murine podocytes (Exo-only, LNA-21-only, Exo-LNA-21) stained for F-Actin (white), FAM-LNA-21 (green), CD9 (magenta) and Nuclei (blue) with merged channels. Scale bars = 20 μm.

**Supplementary Figure 2.**
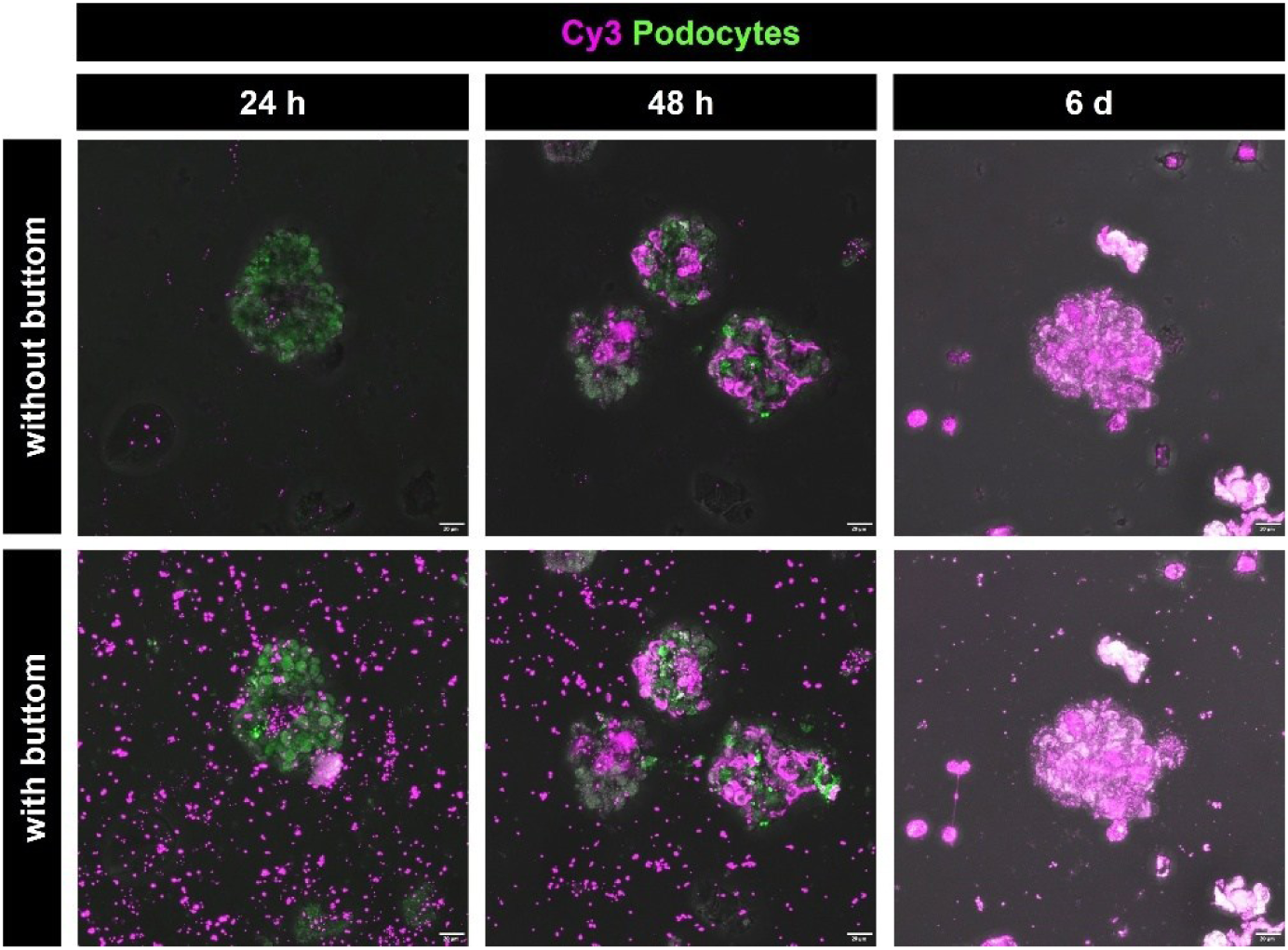
Exosomal Anti-miR-Ctrl. uptake by isolated glomeruli under healthy conditions with and without buttom. cLSM time series of glomeruli treated with Exo-Anti-miR-Ctrl. (Cy3: magenta; Podocytes CFP: green; brightfield overlay) at 24 h, 48 h and 6 d. Scale bars = 20 μm.

